# Complementary cortical and thalamic contributions to cell-type-specific striatal activity dynamics during movement

**DOI:** 10.1101/2025.07.11.664473

**Authors:** Enida Gjoni, Ram Dyuthi Sristi, Haixin Liu, Shahar Dror, Xinlei Lin, Keelin O’Neil, Oscar M. Arroyo, Sun Woo Hong, Hannah Kim, Jeffrey Liu, Sonja Blumenstock, Byungkook Lim, Gal Mishne, Takaki Komiyama

**Affiliations:** Dept. of Neurobiology, Center for Neural Circuits and Behavior, Dept. of Neurosciences, University of California San Diego, La Jolla CA USA; Dept. of Electrical and Computer Engineering, University of California San Diego, La Jolla CA USA; Dept. of Computer Science and Engineering, University of California San Diego, La Jolla CA USA; Halicioğlu Data Science Institute, University of California San Diego, La Jolla CA USA

**Author notes:** co-correspondence. co-first authors (Enida Gjoni, Ram Dyuthi Sristi, Haixin Liu).

## Abstract

Coordinated motor behavior emerges from information flow across brain regions. How long-range inputs drive cell-type-specific activity within motor circuits remains unclear. The dorsolateral striatum (DLS) contains direct- and indirect-pathway medium spiny neurons (dMSNs and iMSNs) with distinct roles in movement control. In mice performing skilled locomotion, we recorded from dMSNs, iMSNs, and their cortical and thalamic inputs identified by monosynaptic rabies tracing. An RNN classifier and clustering analysis revealed functionally heterogeneous subpopulations in each population, with dMSNs preferentially activated at movement onset and offset, and iMSNs during execution. Cortical and thalamic inputs were preferentially activated during onset/offset and execution, respectively, though dMSN- and iMSN-projecting neurons in each region showed similar patterns. Locomotion phase-specific rhythmic activity was found in a subset of thalamic dMSN-projecting neurons and dMSNs. Cortex or thalamus inactivation reduced MSN activity. These findings suggest that corticostriatal and thalamostriatal inputs convey complementary motor signals via shared and cell-type-specific pathways.

## Introduction

Generating appropriate motor outputs to effectively interact with the outside world is a major goal of the central nervous system. Recent advances in large-scale recordings have highlighted that many interconnected areas distributed throughout the brain have movement-related activity (*1–5*). Yet the principles that govern information flow among brain areas, such as the function and specificity of long-range connections, are only beginning to be understood (*6–12*). Individual brain areas are themselves composed of diverse cell types, defined by their transcriptional and functional properties as well as anatomical connectivity (*13–17*). Behavior-relevant information encoded in a given presynaptic area may be passed down uniformly across cell types or separately onto specific neuronal subpopulations of their postsynaptic targets. Characterization of neural circuits at this level of specificity is important for a deeper understanding of how coordinated activity among motor brain areas is achieved and how it relates precisely to motor actions.

Within the cortico-basal-ganglia-thalamic loop involved in movement generation and execution (*18–21*), the dorsolateral striatum (DLS) integrates excitatory inputs from both cortex and thalamus. Striatal direct- and indirect-pathway medium spiny neurons (dMSNs and iMSNs) send distinct axonal projections to the basal ganglia output nuclei via the direct or indirect pathways and have been associated with different roles during movement execution. It is believed that dMSNs play a role in the initiation and facilitation of selected motor sequences, whereas iMSNs are involved in switching between different sequences or inhibiting unwanted motor programs (*22–26*). Previous studies have shown that both dMSNs and iMSNs are active during movement, but they exhibit distinct activity patterns (*27–32*). Whether these differences are inherited from their upstream inputs or shaped locally within the DLS is unknown.

In particular, among the cortical regions that project to the dorsolateral striatum (*33–35*), primary and secondary (M1 and M2) motor cortices are critically involved in motor preparation, execution, and learning (*36–41*). Among the thalamic nuclei, the parafascicular nucleus of the thalamus (PF) represents the major input to MSNs (*33, 35, 42*) and participates in movement execution (*43–45*).

It is unknown how cortical and thalamic neurons contribute to the cell-type-specific activity of dMSNs and iMSNs, especially given that the degree of overlap in their connections with dMSNs and iMSNs is debated (*46, 47*).

We investigated the activity dynamics of dMSNs and iMSNs, as well as their cortical and thalamic inputs, by combining monosynaptically restricted trans-synaptic rabies virus with conditional gene expression together with 2-photon imaging, as mice performed a motor task. We developed and applied a recurrent neural network (RNN) classifier on their activity to determine the degree of functional differences between the two striatal cell types and between their cortical and thalamic inputs. We identified heterogeneous functional subpopulations within MSNs and within their inputs, aligned with specific movement phases. Many dMSNs showed activity around movement initiation and termination, while more iMSNs were active during the movement. The activity of their cortical and thalamic inputs exhibited contrasting dynamics, suggesting a complementary contribution of these inputs to striatal activity. However, within each input region, dMSN-projecting and iMSN-projecting populations showed similar activity patterns. Notably, activity related to forelimb positions was predominantly found in thalamocortical neurons, especially those projecting to dMSNs, suggesting that kinematic information might be conveyed to MSNs in a cell-type-specific manner.

## Results

### Population activity patterns of dMSNs and iMSNs and their cortical and thalamic inputs

To characterize the activity patterns of dMSNs and iMSNs in the DLS, and of the neuronal subpopulations within the cortex and thalamus that project onto them, we conducted *in vivo* two-photon imaging in head-fixed mice while they walked on a motorized circular ladder. In each trial, an auditory cue preceded ladder rotation at a constant speed. (Fig.1A). To selectively label and image the activity of dMSNs and iMSNs, a Cre-dependent viral vector of GCaMP6f was injected in the DLS of specific Cre lines (Drd1-Cre for dMSNs, Adora2a-Cre for iMSNs). To record the activity of cortical or thalamic neurons that project to dMSNs or iMSNs, Cre-dependent helper viruses that allow infection and trans-synaptic spread of monosynaptic pseudo-typed rabies virus (*48*) were injected in the DLS of D1R-Cre and A2A-Cre lines (Fig.1B). After 3-4 weeks, EnvA-pseudotyped rabies that expresses GCaMP6f was injected in DLS (Fig. S1). Imaging with rabies virus was limited to 5-9 days from infection (*49*). Cortical neurons were imaged through a chronic glass window, while striatal and thalamic neurons were imaged via a GRIN lens (Fig.1C). Muscimol injection in each input region led to almost complete abolishment of the movement-related striatal activity (Fig. S2, paired Wilcoxon rank-sum test, p<0.001 for all comparisons after vs. before manipulation), confirming that these three inputs are required for driving DLS activity during the ladder task.

**Fig. 1.**
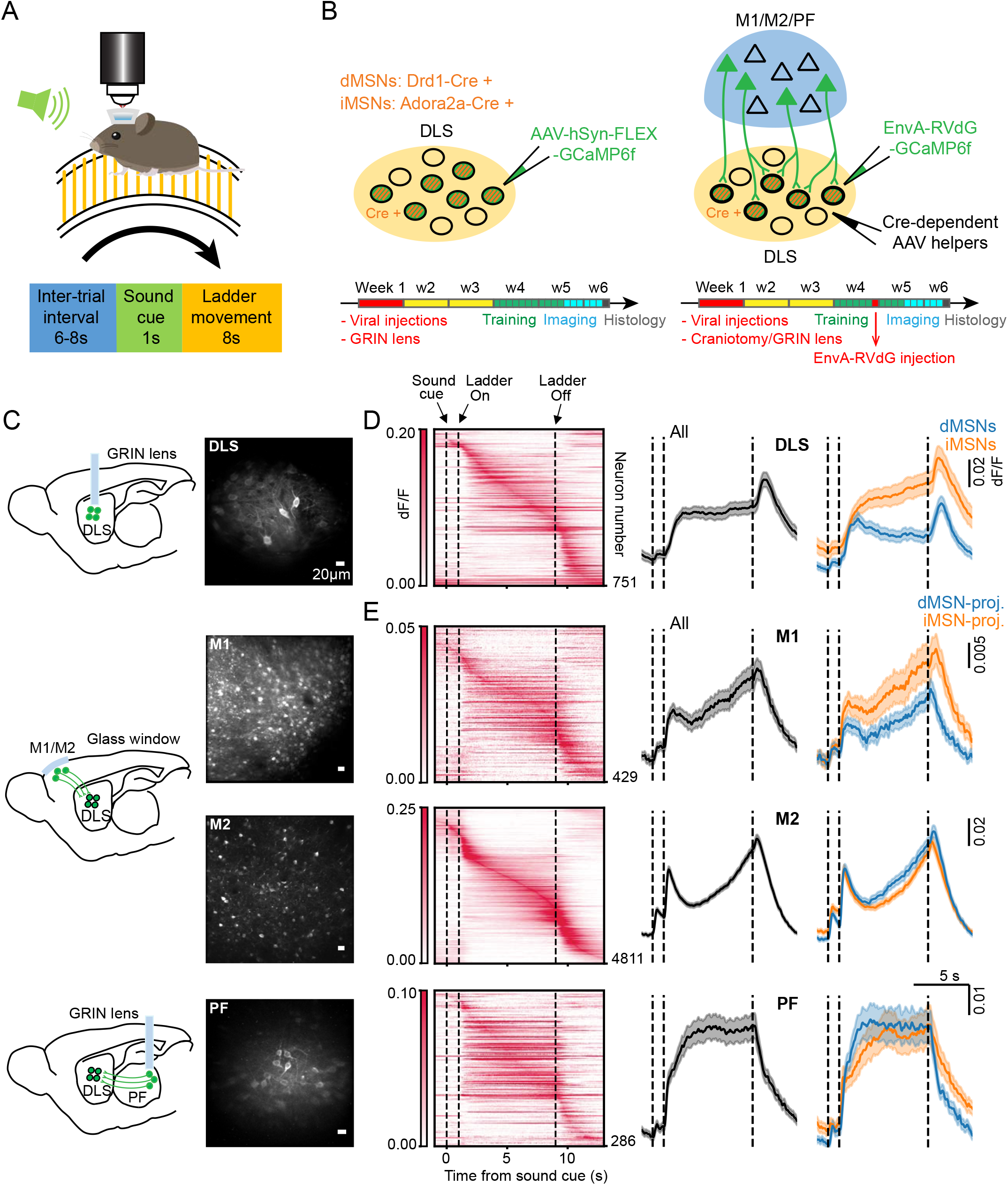
Activity patterns of MSNs and their cortical and thalamic inputs. **A)** Schematic of the motor task. **B)** Left, schematic of the approach used for labeling dMSNs and iMSNs in DLS. Bottom, timeline of experiments. Right, schematic of the approach used for labeling the presynaptic cortical and thalamic inputs to dMSNs and iMSNs. **C)** Schematic of the imaged brain regions and maximum intensity projection images from 2-photon in vivo imaging, showing neurons that express GCaMP6f in DLS, M1, M2, and PF. Scale bar: 20µm. **D)** Left, trial-averaged activity heatmaps of neurons in DLS, ordered based on the maximum peak time of their activity. Middle, population average activity of all DLS neurons. Right, population average activity of dMSNs (blue) and iMSNs (orange), mean ± SEM. n = 337 dMSNs (from N = 21 mice) and n = 414 iMSNs (from N = 12 mice). **E)** Same as D) for the input neurons in M1, M2 and PF. M1: n = 180 dMSN-projecting neurons (from N = 5 mice) and 249 iMSN-projecting neurons (from N = 4 mice); M2: n = 1849 dMSN-projecting neurons (from N =12 mice) and 2962 iMSN-projecting neurons (from N = 12 mice); PF: n = 136 dMSN-projecting neurons (from N = 8 mice) and 150 iMSN-projecting neurons (from N = 9 mice). DLS: dorsolateral striatum; M1: primary motor cortex; M2: secondary motor cortex; PF: parafascicular nucleus of the thalamus.

The average population neuronal activity differed between the four regions (Fig. 1D, E). PF neurons were strongly active throughout the movement period. In contrast, M1 and M2 neurons showed more pronounced activity around the onset and offset of ladder movement. MSN activity appeared to be a combination of the CTX and PF patterns, with both sustained activity during movement as well as activity at ladder onset and offset. These results suggest a complementary contribution of cortical and thalamic inputs to striatal MSN activity during movements.

The population average activity of dMSNs and iMSNs exhibited different patterns. dMSN activity peaked at the ladder onset and declined during the movement before peaking again at the ladder offset, while iMSNs exhibited a consistent increase in activity during movement. In the input areas, the population average activity patterns among dMSN- and iMSN-projecting were similar.

### Cell-type classification based on ensembles of single trial neuronal activity

To explore how different the activity of single neurons is between dMSNs and iMSNs and between dMSN- and iMSN-projecting neurons in each input region, we trained a classifier that predicts the cell type based on the neuron’s activity. We designed the Trial Ensemble Attention Network (TEA-net) (Fig. 2A, top) to capture the nonlinearity and temporal nature of neuronal activity, as well as the variability of a neuron’s activity across single trials in a fixed-length task. Our deep network model is based on an encoder-decoder framework using recurrent neural networks (RNNs) with an “attention”-based mechanism (*50*). The input to the classifier is the neural activity (from 2 sec before to 4 sec after the ladder movement period) in an ensemble of a fixed number of randomly selected trials (Fig. 2A, bottom). This ensemble activity is mapped to a latent representation, which is then used to predict the cell type. The classifier was trained using 10-fold cross-validation across sessions, such that each session was used in the test set once for each neuron. Using the TEA-net classifier trained on ensembles of trial activity we found that dMSNs and iMSNs, as well as dMSN- and iMSN-projecting neurons, are distinguishable above chance level, although the classification performance was generally modest (Fig. 2C, Wilcoxon rank-sum test, p<0.001 for all comparisons; accuracy (%), mean across cell types ± SD: DLS, 63.21 ± 3.98; M1, 62.87 ± 3.43; M2, 59.25 ± 2.06; PF, 64.99 ± 6.94). The classification performance was not statistically significant between the four regions, suggesting a similar degree of distinction between target-cell types and between their input-cell types in motor cortex and thalamus (Wilcoxon rank-sum test with post-hoc Bonferroni correction for n = 6 comparisons: DLS-M1, raw p = 0.791, p adj. = 4.748; DLS-M2, raw p = 0.021, p adj. = 0.127; DLS-PF, raw p = 0.733, p adj. = 4.402; M1-M2, raw p = 0.011, p adj. = 0.068; M1-PF, raw p = 0.571, p adj. = 3.425; M2-PF, raw p = 0.162, p adj. = 0.972). Our TEA-net classifier outperformed other linear classifiers (SVM) and nonlinear fully connected neural networks (FCNN) (Table S1). We then examined the mean activity of correctly classified neurons (Fig. 2D, E). As expected, the activity of these correctly classified neurons showed more differences across cell types (Fig.1D-E). In DLS, correctly classified iMSNs showed higher activity levels that culminated at the end of movement compared to dMSNs (Fig. 2D). Correctly classified iMSN-projecting M1 neurons also showed overall higher activity levels throughout movement (Fig. 2E), whereas correctly classified dMSN-projecting neurons in both M2 and PF showed stronger movement-related activity. The mean activity of the misclassified neurons resembled the activity pattern of the correctly classified neurons of the opposite cell-type. These results indicate that the activity of cell types in each region is somewhat distinguishable, but substantial fractions of them display activity patterns typical of the other cell type.

**Fig. 2.**
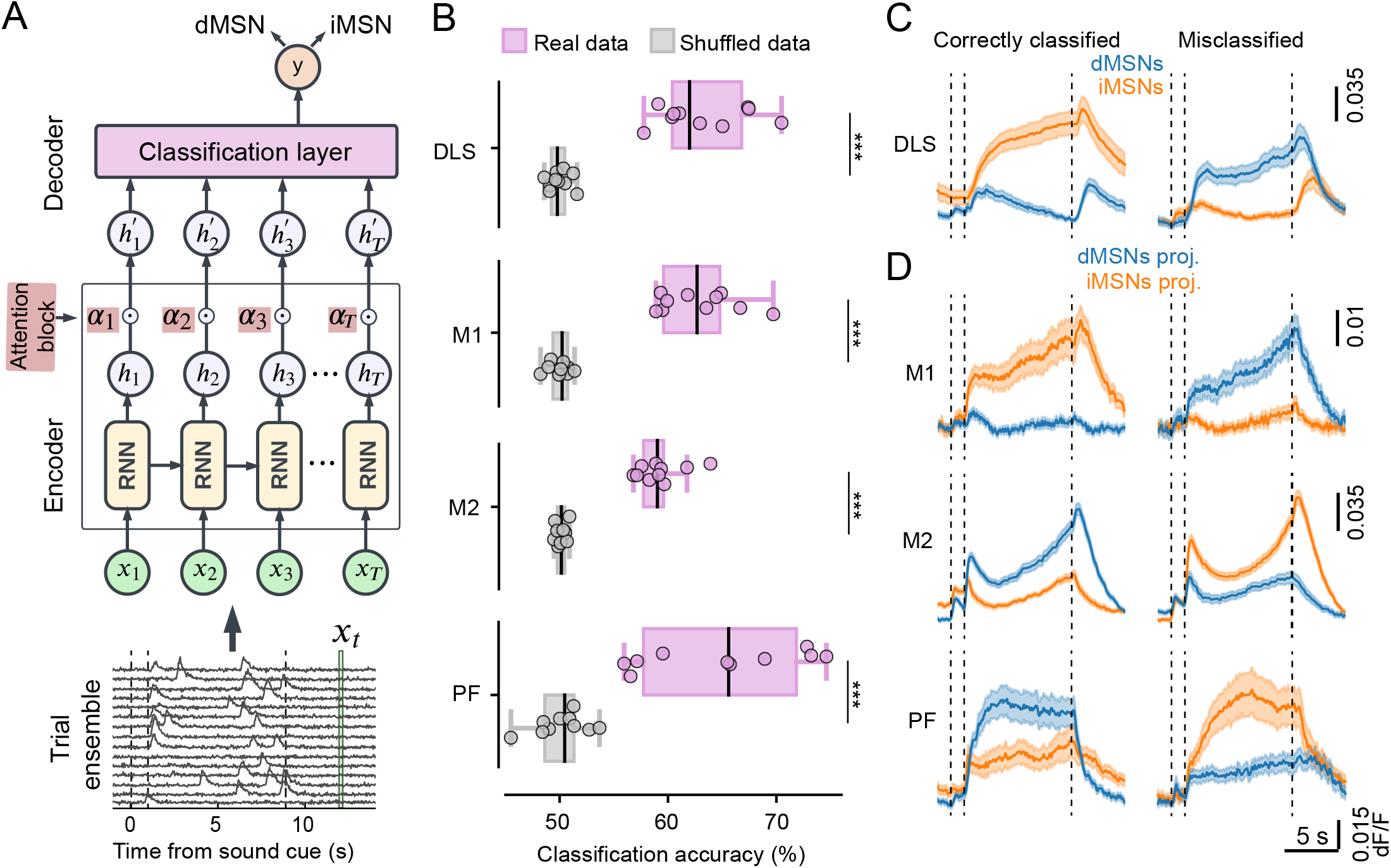
Cell-type classification based on ensembles of single trial neuronal activity. **A)** Top, schematic of the TEA-net classifier. The encoder consists of RNN units that learn a hidden state multivariate time series representation ***h***_***t***_, these are pointwise-weighted by attention values *α*_***t***_ (a learned function of ***h***_***t***_) and then summed to produce 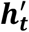 the final latent space representation. This is fed into the decoder, a fully connected classification layer. Bottom, classifier input is based on subsampled ensembles of a fixed number of trials. Each trial includes neural activity from 2 sec before to 4 sec after the ladder movement period. ***x***_***t***_ corresponds to neural activity at time *t* for all the trials in the ensemble. **B)** Cell-type classification accuracy is above chance in each region. Wilcoxon rank-sum test, p<0.001 for all comparisons vs. shuffled data. The classification performance was not statistically different between the four regions. Wilcoxon rank-sum test with post-hoc Bonferroni correction for n = 6 comparisons: DLS-M1, raw p = 0.791, p adj. = 4.748; DLS-M2, raw p = 0.021, p adj. = 0.127; DLS-PF, raw p = 0.733, p adj. = 4.402; M1-M2, raw p = 0.011, p adj. = 0.068; M1-PF, raw p = 0.571, p adj. = 3.425; M2-PF, raw p = 0.162, p adj. = 0.972. **C)** Average activity of correctly classified and misclassified neurons in DLS (mean ± SEM). Total number of correctly classified neurons = 474 (214 dMSNs and 260 iMSNs) and total number of misclassified neurons = 277 (123 dMSNs and 154 iMSNs). **D)** Average activity of correctly classified and misclassified neurons in M1, M2, and PF. M1: total number of correctly classified neurons = 267 (110 dMSN-projecting and 157 iMSN-projecting neurons); total number of misclassified neurons = 162 (70 dMSN-projecting and 92 iMSN-projecting neurons). M2: total number of correctly classified neurons = 2848, of which n = 1100 dMSN-projecting and n = 1748 iMSN-projecting neurons; total number of misclassified neurons = 1963 (749 dMSN-projecting and 1214 iMSN-projecting neurons). PF: total number of correctly classified neurons = 184 (92 dMSN-projecting and 92 iMSN-projecting neurons); total number of misclassified neurons = 102 (44 dMSN-projecting and 58 iMSN-projecting neurons).

### Heterogeneous and distinctive activity patterns in DLS and input areas

To explore the heterogeneity of activity patterns and identify those that are distinctive of each cell type, we performed hierarchical clustering of the latent space representations of the TEA-net model (i.e. the inputs to the classifier). We used the latent space representations as the classifier has been trained to capture separation between cell-types, and this representation has a higher tendency to cluster than raw activity (Fig. S3A). We applied a Dynamic Tree Cut algorithm (*51*) to the hierarchical clustering dendrogram together with subsampling (Methods, Fig. S3B, C) to determine the optimal number of clusters.

For each neuron that belongs to a specific latent space representation cluster, we plotted the corresponding trial-averaged neuronal activity (Fig. 3). We identified several major clusters in each region indicating that functionally heterogeneous subpopulations exist (Fig. 3A-D). Examining the classifier performance of neurons belonging to each cluster, we found that some clusters showed much higher classification accuracies than the population mean accuracy (Fig. 3E). In many of these clusters with high classification accuracy, one of the two cell-types dominated the cluster, indicating that these activity patterns were found primarily in one of the two cell-types (e.g., dMSN-dominant, DLS clusters 1 and 2; iMSN-dominant, DLS clusters 4 and 5). However, in other clusters with high classification accuracies, cell-type dominance was not apparent (e.g., PF clusters 2 and 4) (Fig. S3D), suggesting that these clusters are further heterogeneous within them and additional latent single-trial features might contribute to classifier performance.

**Fig. 3.**
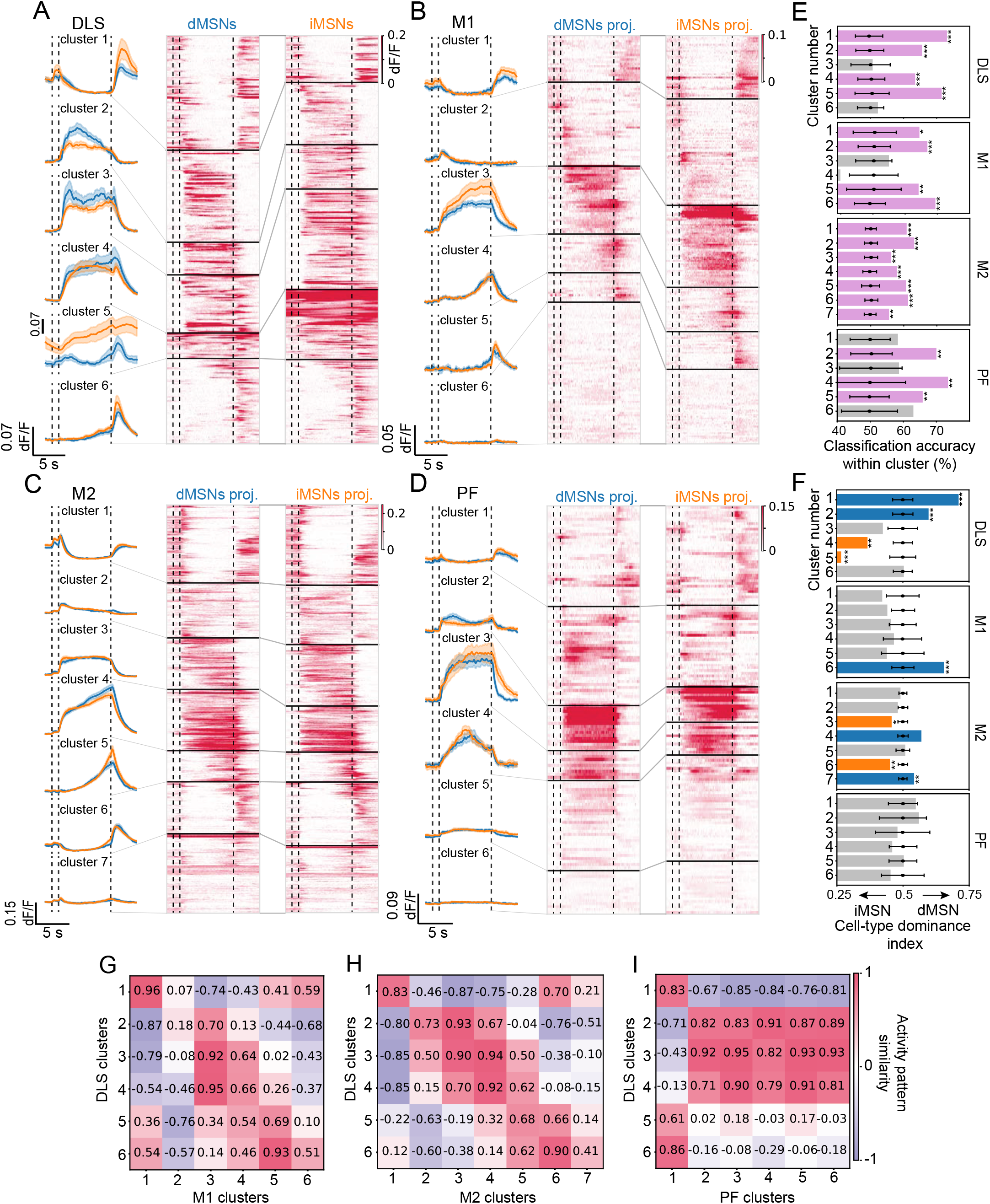
Heterogeneous and distinctive activity patterns in DLS and input regions. **A)** Left, population average activity of dMSNs (blue) and iMSNs (orange) that belong to each cluster. Middle and right columns, trial-averaged activity of individual dMSNs and iMSNs, respectively (mean ± SEM). Neuron order in each cluster is according to the leaf sequence returned by the hierarchical clustering dendrogram. n for dMSNs and iMSNs: cluster 1: 94 and 47; cluster 2: 76 and 63; cluster 3: 27 and 45; cluster 4: 48 and 102; cluster 5: 21 and 71; cluster 6: 71 and 86. **B-D)** Same as A) but for dMSN- and iMSN-projecting neurons in M1, M2, and PF. n for dMSN-projecting and iMSN-projecting neurons: M1, cluster 1: 20 and 38; cluster 2: 37 and 65; cluster 3: 30 and 50; cluster 4: 17 and 27; cluster 5: 13 and 23; cluster 6: 63 and 46; M2, cluster 1: 352 and 585; cluster 2: 249 and 431; cluster 3: 234 and 445; cluster 4: 277 and 335; cluster 5: 141, and 218; cluster 6: 236 and 461; cluster 7: 360 and 487; PF, cluster 1: 33 and 36; cluster 2: 33 and 30; cluster 3: 15 and 13; cluster 4: 10 and 12; cluster 5: 30 and 39; cluster 6: 15 and 20. Dashed lines from left to right in each plot indicate: sound cue onset, ladder onset, and ladder offset. **E)** Classification accuracy between cell types in each cluster, for each region. Black circles and corresponding error bars indicate mean ± SD of the chance accuracy obtained by shuffle control. Bars in purple indicate cluster accuracy values that were statistically higher than the shuffled distribution. * p < 0.05, ** p < 0.01, *** p < 0.001. **F)** Cell-type dominance in each cluster, defined as the ratio between the normalized fraction of one cell type and the normalized fraction of both cell types. Values above 0.5 indicate dMSN (or dMNS-proj.) dominance, whereas values below 0.5 indicate iMSN (or iMSN-proj.) dominance. Black circles and corresponding error bars indicate mean ± SD of the chance cell-type dominance obtained by shuffle control. Bars in blue and orange indicate significantly dMSN-(or dMNS-proj.) and iMSN- (or iMSN-proj.) dominant clusters, respectively. * p < 0.05, ** p < 0.01, *** p < 0.001. **G-I)** Similarity matrices built by computing the Pearson’s correlation coefficients between average neuronal responses of DLS clusters and M1, M2, PF clusters.

DLS clusters enriched in dMSNs showed strong activity near the onset and offset of movement (Fig. 3A, F, clusters 1 and 2) while iMSN-dominant clusters (clusters 4-5) exhibited peak activity toward the later phase of movement. In M1 and M2, many clusters were formed by neurons that were active around the onset or offset of the movement (Fig. 3B-C, M1 clusters 1, 2, 4 and 5; M2 clusters 1-2 and 5-7). Additionally, in M2, about a third of the neurons belonged to movement-active clusters, with the cluster with consistent activity throughout the movement containing more iMSN-projecting neurons (Cluster 3), and the cluster with activity ramping towards the end of the movement dominated by dMSN-projecting neurons (Cluster 4). Furthermore, a larger fraction of dMSN-projecting M1 and M2 neurons did not show clear activity time-locked to our task (M1 cluster 6 and M2 cluster 7), raising the possibility that these neurons may relay information selectively to dMSNs in behavioral contexts that are not explored in our study. In contrast to cortical inputs, the majority of PF clusters showed activity during the movement (Fig. 3D, clusters 2-5).

The results indicate that dMSNs and iMSNs and their projecting neurons display specific movement-related activity patterns, some of which are more common in one cell-type than the other. We assessed the similarity between DLS clusters and input neuron clusters (Fig. 3H-J) by computing Pearson’s correlation between their cluster-averaged activity patterns. M1 and M2 clusters more closely resembled movement onset/offset clusters in DLS, as compared to PF. The M1 movement cluster was more similar to late-movement clusters in DLS, while all PF and M2 movement clusters were highly similar to all DLS movement clusters. These results suggest that corticostriatal neurons might play an important role in driving striatal activity at movement start and end, which are more prominent in dMSNs. Thalamostriatal neurons may be more important in driving striatal activity during movements.

### Forelimb-related rhythmicity is input- and cell-type specific

To explore how the neural activity of the two MSN cell types and their specific cortical and thalamic inputs relates to limb kinematics in this motor task, we extracted the time series of the coordinates of the four paws using DeepLabCut (*52*) (Fig. 4A). To detect neurons that show rhythmic activity with similar periodicity as the contralateral left forepaw during locomotion, we performed a Fourier analysis on the left forelimb trajectory and on neural activity during the movement period of each trial (Methods, Fig. 4B). We found that a small fraction of DLS neurons showed activity matching the rhythmicity of the forelimb, with a higher fraction of rhythmic cells found in dMSNs than in iMSNs (Fig. 4C, dMSN rhythmic cells n = 17 out of 308 (5.5%), iMSN rhythmic cells n = 6 out of 400 (1.5%); Fisher exact test p raw = 0.004, with Bonferroni-adjusted p = 0.018). Among the input regions, very few rhythmic cells were observed in M1 and M2 (Fig. 4D, M1: dMSN-proj. rhythmic cells n = 2 out of 166 (1.2%), iMSN-proj. rhythmic cells n = 5 out of 234 (2.1%); M2: dMSN-proj. rhythmic cells n = 14 out of 1881 (0.7%), iMSN-proj. rhythmic cells n = 20 out of 2865 (0.7%)). In contrast, we found a substantial proportion of rhythmic cells in PF neurons, with dMSN-projecting thalamic neurons containing a significantly higher fraction of rhythmic cells (dMSN-proj. rhythmic cells n = 58 out of 202 (28.7%), iMSN-proj. rhythmic cells n = 19 out of 160 (11.9%); Fisher exact test p raw < 0.001, with Bonferroni-adjusted p< 0.001).

**Fig. 4.**
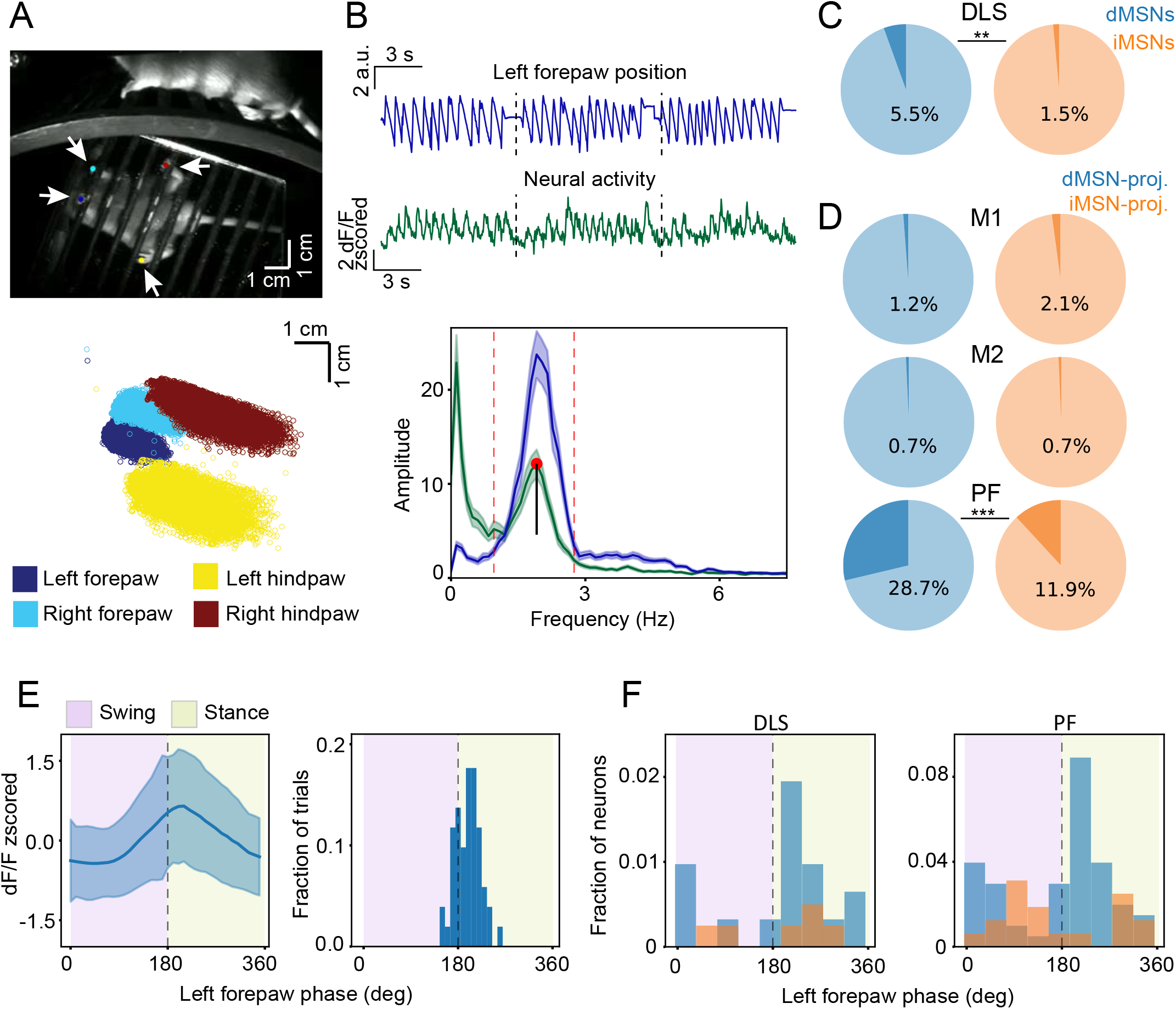
Forelimb-related rhythmicity is input- and cell-type specific. **A)** Top, example video frame of a mouse on the motorized ladder and the marked limbs (mirror view) used for training DeepLabCut. Bottom, coordinates of the tracked limbs of a mouse during one imaging session. **B)** Top, in dark blue, example trace of the first PCA component of the *x* coordinate of the left forelimb during three consecutive trials in one imaging session (from ladder onset to 1 s after the ladder offset). In green, corresponding activity trace of a PF neuron recorded during the same imaging session as above. Note that neural activity follows the periodicity of the forelimb position. Bottom, power spectral densities of the forelimb trajectory and neural activity, averaged across all trials of the same imaging session as above. The dotted lines indicate the full width at 10% maximum of the peak frequency of the forelimb trajectory. Neural activity that shows local maximum peak (red dot) within this frequency interval and peak prominence (black line) above a defined threshold is defined as ‘forelimb-related rhythmic’. **C)** Fraction of dMSNs and iMSNs with forelimb-related rhythmic activity (dMSN rhythmic neurons = 17 out of 308, iMSN rhythmic neurons = 6 out of 400). Fisher exact test p raw = 0.004, with Bonferroni-adjusted p = 0.018. **D)** Fraction of dMSN- and iMSN-projecting neurons with forelimb-related rhythmic activity in M1, M2 and PF. M1: dMSN-projecting rhythmic neurons = 2 out of 166, iMSN-projecting rhythmic neurons = 5 out of 234, Fisher exact test p raw = 0.704, with Bonferroni-adjusted p = 1; M2: dMSN-projecting rhythmic neurons= 14 out of 1881, iMSN-projecting rhythmic neurons = 20 out of 2865, p raw = 0.862, with Bonferroni-adjusted p = 1; PF: dMSN-projecting rhythmic neurons = 58 out of 202, iMSN-projecting rhythmic neurons = 19 out of 160; p raw < 0.001, with Bonferroni-adjusted p<0.001. Comparisons between regions: Fisher exact test, DLS vs. M1, p raw = 0.178, with Bonferroni-adjusted p = 1; DLS vs. M2, p raw < 0.001, with Bonferroni-adjusted p < 0.001; DLS vs. PF, p raw < 0.001, with Bonferroni-adjusted p < 0.001; PF vs. M1, p raw < 0.001, with Bonferroni-adjusted p < 0.001; PF vs. M2, p raw < 0.001, with Bonferroni-adjusted p < 0.001; M1 vs. M2, p raw = 0.037, with Bonferroni-adjusted p = 0.219; **E)** Left, trial-averaged activity (mean ± SD) of an example dMSN-projecting neuron in PF with forelimb-related rhythmic activity that peaks at the beginning of the stance phase. Right, histogram of the neural activity peak relative to the left forelimb phase, at each trial for the same neuron (n = 51 trials). **F)** Left, histogram of the forelimb-position phase tuning of dMSNs and iMSNs in DLS with forelimb-related rhythmicity. Right, same for dMSN- and iMSN-projecting neurons in PF. Phase-tuning comparisons: dMSN-vs. iMSN-projecting in PF, Watson two-sample U2 test, p = 0.011; dMSNs in DLS vs. dMSN-projecting in PF, Watson two-sample U2 test, p = 0.696.

Each stride in the gait cycle consists of a stance phase, when the forelimb is in contact with the ladder bar, followed by a swing phase, when the forelimb is raised and pushed forward. We examined whether rhythmic neurons are preferentially tuned to a specific phase of left forelimb position (Fig. 4E). We found that thalamic dMSN- and iMSN-projecting neurons exhibited opposite phase-tuning, with dMSN-projecting neuron activity peaking during the stance phase and iMSN-projecting neuron activity peaking during the swing phase (Fig. 4F, Watson two-sample U^2^ test, p=0.011). dMSNs in DLS showed a similar forelimb phase preference to thalamic dMSN-projecting neurons. These findings suggest that the contralateral forelimb position during movement is primarily encoded by thalamostriatal neurons, rather than by corticostriatal neurons. Furthermore, thalamostriatal neurons may provide information related to forelimb position to striatal neurons via target cell-type specific connections.

## Discussion

This study characterized the activity dynamics and kinematic-related information of pre- and postsynaptic populations of four motor areas. By imaging DLS cell types and their cortical and thalamic presynaptic inputs, we aimed to identify cell-type specific differences of activity and whether they arise from specificity of presynaptic inputs.

We identified subpopulations with distinct activity patterns within MSNs and within their cortical and thalamic inputs. All neuronal groups (dMSNs, iMSNs, and their respective inputs) showed heterogeneous activity patterns and we did not find any activity pattern that was only observed in one cell type. Nevertheless, dMSNs were predominant in clusters with activity around movement initiation and termination, while more iMSNs were active during movements. This finding aligns with previous studies that have highlighted co-activation of the two cell types but with distinct patterns (*27–32*). The majority in corticostriatal and thalamostriatal populations showed activity around onset/offset of movement and during movement, respectively. This suggests a complementary contribution of these inputs to striatal activity, with cortical and thalamic activity more closely matching that of dMSNs and iMSNs, respectively. Indeed, even though cortical and thalamic axons innervate both dMSNs and iMSNs, it has been suggested that cortical and thalamic inputs are biased towards dMSNs and iMSNs, respectively (*46, 53–56*), although an opposite observation has also been made for the thalamostriatal projections (*54*). These observations together suggest that corticostriatal and thalamostriatal connections have distinct functions, with corticostriatal neurons driving striatal activity primarily at the onset and offset of movements, while thalamostriatal neurons contribute more to striatal activity during the movements. These findings suggest that striatal activity during movements arises as an integration of both corticostriatal and thalamostriatal inputs, instead of being primarily driven by cortical activity (*57, 58*).

Our clustering analysis did not reveal clear differences in dMSN- and or iMSN-projecting neurons that correspond to differences in dMSNs and iMSNs. However, in a separate analysis, we found movement kinematics-related neuronal rhythmicity primarily in PF but also in DLS. Importantly, a majority of these neurons in PF were dMSN-projecting neurons, and a majority in DLS were dMSNs with phase tuning matching that of dMSN-projecting PF neurons. Taken together, the findings suggest that while dMSN- and iMSN-projecting neurons in each region might share many activity patterns (and some are likely overlapping populations), a subset of them encode and transmit kinematic or task-related information in a cell-type specific manner. Future studies should further test this idea with simultaneous recordings of identified pre- and post-synaptic populations as well as pathway- and cell-type-specific activity manipulations. Furthermore, kinematic encoding in the PF-DLS pathway should be further studied to determine if it is restricted to rhythmic locomotion studied here or more broadly observed in other movement contexts. We note that studies with electrophysiological recordings have described a higher fraction of locomotion-related rhythmic cells both in the DLS as well as in M1 compared to what we report here (*59, 60*). Our study using calcium imaging may be underestimating rhythmic activity.

Machine learning-based methods are becoming increasingly popular for identifying cell types based on neuronal activity (*61–64*). In this study, we used a supervised classification approach based on recurrent neural networks as a tool for determining functional differences among cell types. Our classifier design combined trial ensemble inputs and nonlinear temporal modeling via the RNN, and outperformed baseline models under cross-validation. By incorporating neuronal activity across multiple single trials, rather than using trial-averaged activity the classifier may capture features—such as fraction of responsive trials, response amplitude and timing, and their consistency across trials—that determine how reliably and strongly information is encoded and transmitted within circuits. These features may prove crucial for both determining specificity and connectivity rules in cell-type defined circuits. While our design accounts for the nonlinearity of encoding, temporal structure of neuronal activity, and single-trial activity, it is aligned to task events and does not account for trial-by-trial variability in the movements of the animal. Incorporating kinematic features, e.g., paw position and velocity, in future designs would have the potential to enhance the quantification of functional diversity of neuronal populations in relation to the generation of movements and offer mechanistic insights into how cell-type specific circuits generate precise movements.

### Experimental methods Animals

All animal procedures were performed in accordance with guidelines and protocols approved by the UCSD Institutional Animal Care and Use Committee and the National Institutes of Health (NIH). Mice were group-housed in disposable cages with standard bedding in a temperature-controlled room (∼21 °C) with a reversed light cycle (10.00–22.00 corresponded to the dark period). Mice were allowed *ad libitum* access to food and water. Experiments were performed in two transgenic mouse-lines: B6.FVB(Cg)-Tg(Adora2a-Cre)KG139Gsat/Mmucd (RRID:MMRRC_036158-UCD) and B6.FVB(Cg)-Tg(Drd1-Cre)FK150Gsat/Mmucd (RRID:MMRRC_036916-UCD). Both males and females were used for experiments. All experiments were performed during the dark period.

### Surgeries and virus injections

For all surgeries, adult mice (6-10 weeks old) were anesthetized with 1-2% isoflurane and injected with Baytril (10 mg/kg), dexamethasone (2 mg/kg), and buprenorphine (0.1 mg/kg) subcutaneously at the beginning to prevent infection, inflammation, and pain. Recovery was monitored daily for 3 consecutive days following surgery with administration of Baytril, dexamethasone, and buprenorphine.

For 2-photon imaging of dMSNs and iMSNs in the dorsolateral striatum, AAV1-hSyn-FLEX-GCaMP6f (Addgene, Catalog # 100833-AAV1) was injected via beveled glass pipettes into the right hemisphere of Drd1-Cre and Adora2a-Cre mice respectively, at coordinates: 0.5 mm anterior and 2.2 mm lateral from bregma, at depths 2.2, 2.3 and 2.4 mm from the pia (∼200 nL at each depth, at a rate of 20 nL per minute. The pipette was kept in the brain for ∼15 minutes to avoid virus leakage. For GRIN lens implantation in DLS (or PF, see below), a similar procedure to (*65*) was followed. GRIN lenses of 0.5 or 0.6 mm diameter (Inscopix, product number 1050-006232 and 1050-004597) were lowered into the above-mentioned coordinates at a depth of 2 mm for DLS imaging. GRIN lens implantation was performed within a week from viral injection. Imaging was usually performed 3-4 weeks after lens implantation.

For 2-photon imaging of the cortical or thalamic inputs that project onto dMSNs or iMSNs in the dorsolateral striatum, a 1:1 or 1:2 solution mix of helper viruses AAV-EF1a-DIO-mRuby2-P2A-TVA and AAV-EF1a-DIO-oPBG was injected into the right hemisphere of Drd1-Cre and Adora2a-Cre mice respectively, at coordinates: 0.5 mm anterior and 2.2 mm lateral from bregma, at depths 2.2, 2.3 and 2.4 mm from the pia (∼0.2 μL at each depth). In the case of M1 imaging, a 22° angled injection into the DLS was performed in order to avoid inserting the pipette directly through M1, at coordinates: 0.5 mm anterior, 3.17 mm lateral from bregma. On the same day, a craniotomy centered around 0.5 mm anterior and 1.5 mm lateral from bregma or around 2.3 mm anterior and 1.2 mm lateral from bregma, was performed as previously described (*37, 40, 66, 67*) for imaging M1 or M2, respectively. Otherwise for PF imaging, a GRIN lens was lowered into the brain at coordinates: 2.2 mm posterior and 0.65 mm lateral from bregma at a depth of 3.2 mm. After 3-4 weeks, EnvA-pseudotyped rabies that expresses GCaMP6f (EnvA-RVdG-mRuby-GCaMP6f, in the case of M2 imaging or EnvA-RVdG-GCaMP6f in the case of M1 and PF imaging) was injected into DLS at the same coordinates and depths as mentioned above (∼0.2 μL at each depth. At a 22° angle for M1 imaging). After virus injection, the glass pipette was left in place for 20-30 minutes to minimize off-target expression. Imaging with rabies virus was limited to 5-9 days from infection (*49*). AAV helper and EnvA-pseudotyped rabies viruses were produced in Prof. Lim’s laboratory at UCSD.

### Motorized ladder task

Head-fixed mice were positioned on a custom-built, circular ladder (diameter = 18 cm, rung diameter = 0.3 cm, rung spacing = ∼1 cm) motorized by an electric DC motor (12V, 60 RPM). Trial structure was controlled using BPod v0.5 (https://sanworks.github.io/Bpod_Wiki/). In each trial, 8 seconds of ladder rotation (speed ∼ 10 mm/s) followed an auditory cue (1 s at 3 kHz). Trials were interleaved by a variable 6-8 second inter-trial interval. In each trial, the auditory cue (= trial start) followed a ∼100 ms infrared LED light that was placed outside of the mouse’s view and used for aligning trials during video analysis. Mice were trained on the ladder task for 7 to 12 days, daily, till they became able to walk or run on the rungs by engaging all four limbs. For the experiments where Enva-pseudotyped rabies was used for labelling MSN inputs, mice were trained on the task for 4-5 days before the viral injection, and continued to be trained after, and imaged within 9 days from the viral injection. During 2-photon imaging sessions, two orthogonal views of the mouse were captured simultaneously through a mirror mounted at 45° inside the motor task setup, at 30 Hz with an IR-sensitive video camera (DMK 23U618, The Imaging Source) controlled by IC Capture software, version 2.5.

### 2-photon imaging and analysis

#### Imaging data acquisition

2-photon imaging was conducted with a commercial two-photon microscope (Movable objective microscope (MOM), Sutter Instrument, retrofitted with a resonant galvanometer-based scanning system from Thorlabs), 16× objective (Nikon) and a Ti:Sapphire excitation laser at 925 nm excitation light (Mai Tai, Spectra-Physics) controlled by ScanImage (Vidrio Technologies). For imaging of M1, DLS and PF neurons, images (512 × 512 pixels, 0.98 µm/pixel) were recorded at ∼28 Hz for the duration of the behavioral session (∼20 minutes). For M2 imaging, two fields of view at different depths were acquired near-simultaneously at ∼14 Hz. Frame times were recorded and synchronized with behavioral recordings by the Ephus software (controlled by MATLAB). Multiple non-overlapping fields of view were recorded from each animal in consecutive days.

#### Two-photon imaging analysis

Images were first aligned frame-by-frame using a custom MATLAB program to correct for lateral shifts (*68*). ROIs were manually drawn using a custom MATLAB program (*37, 67*). A ring-shaped background-ROI containing neuropil signal was created from the border of each neuronal ROI to a width of 6 pixels. Fluorescence was processed as described (*37*). Briefly, pixels within each ROI were averaged to create a fluorescence time series, and the background fluorescence was subtracted. The time-varying baseline (F) of the fluorescence trace was estimated by iteratively smoothing inactive portions of the trace. The normalized dF/F trace was then calculated, where dF was determined by subtracting the baseline trace from the raw trace and F is the calculated time-varying baseline. In the case of M2 imaging, dF/F was calculated as the ratio between the GCaMP6f and the mRuby signal. Two criteria were applied for including the recorded neurons in all subsequent analyses: 1) session median raw fluorescence signal above 100 a.u. and 2) at least two activity events (see below) per minute present during the imaging session.

### Muscimol inactivation

Muscimol inactivation experiments were performed on a subset of mice implanted with a GRIN lens in DLS. A field of view in DLS was imaged for approximately 15 minutes as mice performed the ladder task. Immediately after the imaging session, mice were anesthetized as described above and 25 nL muscimol hydrobromide (5 μg/μL, Sigma-Aldrich) was injected with a glass pipette unilaterally into the right hemisphere, at these coordinates: M1: 0.5 mm anterior and 1.5 mm lateral from bregma, depth 0.35mm; M2, 2.3 mm anterior and 1.2 mm lateral from bregma, depth 0.35mm; PF: 2.2 mm posterior and 0.65 mm lateral from bregma, depth of 3.3 mm. Mice were allowed to recover in their home cage and the same field of view was re-imaged after ∼30 minutes from the injection.

### Activity events

For quantifying the effect of muscimol inactivation of inputs on striatal activity, activity events were determined as previously described (*37*) and compared with the imaging sessions before the manipulation. Briefly, events detected sharp rises in the fluorescence trace and were defined if the first derivative (velocity) of the fluorescence trace crossed five times the standard deviation of the inactive portions of the fluorescence trace. Trials that contained at least one activity event were defined as ‘active trials’, and the fraction of active trials was compared before and after the manipulation experiment.

### Histology

At the end of the imaging sessions, mice were transcardially perfused with sodium phosphate buffered saline (PBS) followed by 4% paraformaldehyde (PFA, Sigma-Aldrich) in PBS. Brains were extracted, postfixed in 4% PFA overnight at 4ºC and then stored in 30% sucrose in PBS at 4ºC for 2-3 days. Coronal sections (40 µm thick) were obtained using a Leica SM 2000R sliding microtome. Sections were mounted on slides using CC Mount (Sigma-Aldrich) and imaged with an Axiozoom V16 microscope to confirm viral expression and GRIN lens positioning.

### Computational Methods

#### Notations

We use the following notational convention below. Upper case non-bold for fixed scalars *N*, Upper case bold for matrices ***M***, and lower case bold for vectors ***v***.

### Trial Ensemble Attention Network (TEA-net)

#### Data preprocessing and trial-ensemble construction

For every neuron, dF/F traces were segmented into single-trial windows that started 2 sec before the ladder onset and ended 4 sec after ladder offset. For each region we determined *N*, the smallest number of trials recorded in any session for a given region (DLS: *N* = 42, M1: *N* = 47, M2: *N* = 50, PF: *N* = 28). For each neuron we then generated multiple bootstrap ensembles. We randomly sampled *N* trials without replacement and stacked them into a matrix ***X*** ∈ ℝ^***N*** × ***T***^ to form an ensemble of the neuron’s activity of ***T*** timeframes across *N* trials. The number of ensembles used to train TEA-Net was balanced across cell types in each region to ensure equal class priors. In a given region, we define *N*_all_ as the total number of neurons, with *N*_*d*_ dMSNs (or dMSN-projecting) and *N*_*i*_ iMSNs (or iMSN-projecting) where *N*_*d*_ + *N*_*i*_ = *N*_all_. We generated *N*_all_ × 200 ensembles in total, allocating half (*N*_all_ × 100) to each cell type. Thus, every dMSN (or dMSN projecting) neuron contributed 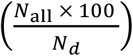 ensembles and every iMSN (or iMSN projecting) neuron contributed 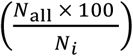 ensembles, ensuring balanced training data for TEA-net.

#### Network architecture

Given an input ensemble ***X***, TEA-net predicts whether its source is a direct or indirect pathway neuron. The network consists of three modules:

1. *Temporal encoder* – a single-layer vanilla recurrent neural network (hidden size = *d*_*h*_= 10) processes the ensemble column-wise, i.e. the input *x*_*t*_ at every timeframe *t* consists of the activity of the neuron at time *t* across the *N* trials. The encoder outputs a hidden state 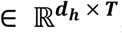, whose *t*-th column is the vector ***h***_***t***_.
2. *Learned temporal attention* – We use a lightweight attention approach, analogous to (*50*), by applying a 1 × 1 convolution across hidden channels to produce a scalar relevance score *e*_*t*_ for every frame. Soft-max normalization yields attention weights *α*_*t*_ that highlight informative hidden representations while down-weighting uninformative ones.
3. *Context aggregation and classification* – the weighted hidden states (*α*_*t*_ · ***h***_***t***_) are summed over the hidden dimension, producing a length-***T*** context vector 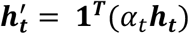, that preserves the temporal nature of the ensemble. A fully connected layer (input = ***T***, output = 2) maps 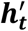 to class logits.

#### Training procedure

Training and evaluation followed a 10-group fold cross-validation at the session level: all neurons recorded in the same imaging session were assigned to the same fold, and every fold contained ∼10 % of the direct- and indirect-pathway neurons. Models were trained for 50 epochs with a batch size of 256 ensembles, an Adam optimizer (*69*) (initial learning rate = 0.001, weight decay = 0.001), and a step-decay scheduler that reduced the learning rate by 90 % every 10 epochs. The loss function was cross-entropy. Early stopping (*70*) (patience = 5 epochs) was applied based on validation loss to prevent over-fitting.

#### Evaluation Metrics

For each test neuron in each cross fold, the majority vote across its 200 ensemble predictions determined the final label. We then compute the average cell type accuracy of the classifier for each region, e.g., for DLS this is given by:

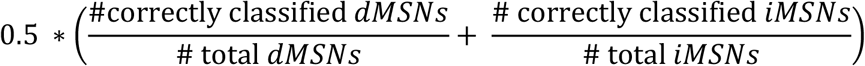

To test whether classifier performance exceeded chance, cell-type labels were randomly reassigned within each session, the entire cross-validation procedure was repeated, and the resulting accuracies were compared to the true-label distribution with a two-sided Mann–Whitney *U*-test.

### Alternative classifier models

We trained two baseline models and evaluated them with the same metrics used for TEA-Net, enabling a direct comparison.

Support-vector-machine (SVM) baselines

As a linear reference, we trained *linear* SVMs (*71*) on two alternative representations of the data:

- Trial-averaged activity: the dF/F trace averaged across all the trials of a neuron, yielding a single ***T***-length vector per neuron. The soft-margin penalty hyper-parameter *C* was tuned by 10-fold cross-validation, searching the grid C ∈ {0.1, 1, 10, 10^2^, 10^3^, 10^4^, 10^5^}; the best score was obtained at *C* = 10.
- Trial-ensemble input: the random ensemble matrix ***X*** flattened into a vector. The same grid search favored a larger penalization constant, *C* = 10^4^.

#### Feed-forward control network with attention (FC-attention)

To test whether TEA-net’s performance benefits from its ability to model temporal dependencies, we built a parameter-matched baseline in which the recurrent temporal encoder is replaced by a time-distributed fully connected linear layer. Concretely, each frame ***x***_***t***_ ∈ ℝ^***N***^ is mapped to a hidden vector ***h***_t_ = ***ReLU***(***Wx***_t_ + ***b***) where 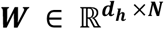, with the same weights applied independently at every *t*. All subsequent stages—learned temporal attention, context aggregation and classification—are identical to those in TEA-net and are applied to the sequence {***h***_***t***_}. Thus, the performance gap quantifies the benefit of the recurrent model (i.e. it is not due to additional capacity or alternative readout mechanisms).

### Hierarchical clustering of latent space trajectories

To uncover functional sub-groups of MSNs and their inputs, we clustered the latent space representation learned by TEA-net for each region. To derive a robust, fold-independent embedding for every neuron, we first averaged over trial ensembles *within* each cross-validation fold: TEA-net projects every bootstrap trial ensemble to a ***T***-dimensional latent trajectory, and the element-wise mean of these trajectories yields a single ***T***-length vector that summarises the fold. Repeating this procedure for all ten folds produces ten such vectors, each residing in the latent space of its respective model. After min–max normalizing each vector within its own fold, we concatenated the ten vectors, resulting in a **10** × ***T*** matrix that serves as the neuron’s final latent representation for subsequent clustering. Pairwise dissimilarities between these vectors were computed with Euclidean distance, and agglomerative hierarchical clustering was performed with Ward’s criterion, which merges clusters to minimize the increase in within-cluster (*72*). Applying Dynamic Tree Cut —specifically the cutreeDynamic function from the R package **dynamicTreeCut** (*51*) —to the resulting dendrogram yielded the cluster assignments.

#### Hyper-parameter selection (deepSplit and minClusterSize)

The two parameters governing dynamic tree cutting are 1) *deepSplit* which governs the *granularity* of cluster division and 2) *minClusterSize* which specifies the size of the smallest allowable cluster as a fraction of the total neuron count. These two parameters were tuned with a 10000-iteration sub-sampling stability analysis. For each brain region, we repeatedly clustered bootstrap subsets containing 80 % of the neurons. We then selected the parameter pair that maximized the fraction of subsets whose cluster count matched the full-data solution, i.e. the mode of the sub-sampling histogram coincided with the cluster number at 100 % sampling. The resulting region-specific settings were: for DLS, *deepSplit*=*3* and *minClusterSize*=0.095; for M1, *deepSplit*=*2* and *minClusterSize*=0.080; for M2, *deepSplit*=*3* and *minClusterSize*=0.070; for PF, *deepSplit*=*3* and *minClusterSize*=0.080.

### Cluster accuracy and cell-type dominance

Two complementary metrics were computed for every cluster returned by the hierarchical procedure: a *normalized classification accuracy* that evaluates how well TEA-net’s cell-type predictions align with ground truth within that cluster, and a *cell-type-dominance index* that quantifies the cluster’s compositional bias toward dMSNs (or dMSN-projecting) or iMSNs (or iMSN-projecting) neurons.

We define the following cell type normalized fractions:

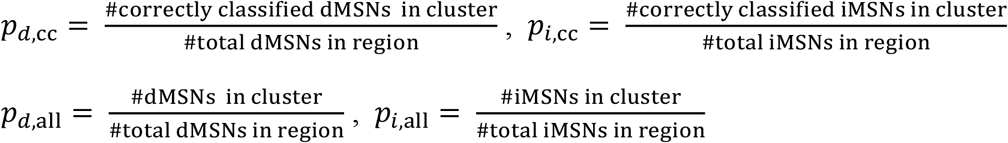

where the subscripts *d* and *i* represent dMSNs and iMSNs (dMSN-projecting and iMSN-projecting for the input regions), respectively, while ‘cc’ indicates correctly classified neurons.

The cluster’s accuracy, adjusted for unbalanced cell types in a given region, was then defined as:

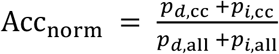

Chance level was obtained by randomly permuting the predicted labels across all neurons (10000 iterations) while keeping cluster membership and total cell-type counts unchanged. One-sided p-values were derived from this null distribution, and clusters with p < 0.05 were deemed statistically significant.

Cell type dominance index was defined as:

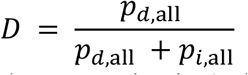

Values above 0.5 indicate dMSN (or dMSN-projecting) dominance, whereas values below 0.5 indicate iMSN (or iMSN-projecting) dominance. Chance level was assessed by shuffling the true cell-type identities across neurons (10000 iterations), recomputing the index, and deriving two-sided p-values.

### Clusterability analysis

To quantify how strongly the data deviates from uniform random distribution, we computed the Hopkins statistic *H* (*73*) for 1) the raw neural activity z-scored after trial-averaging and 2) the normalized TEA-net latent representations used for hierarchical clustering. The test was run with pyclustertend (v 1.3; function pyclustertend.hopkins). For a dataset of n neurons, the routine samples *m* = 0.1 *n* observations—half uniformly random “probe” points drawn from the data hyper-cube and half actual data points. It then compares their nearest-neighbor Euclidean distances, and returns the Hopkins statistic *H*. Values of *H* are interpreted as follows: *H* ≈ 0.5 indicates complete spatial randomness, *H* > 0.5 increasing clusterability, and *H* < 0.5 regular (over-dispersed) structure. For each brain region we generated 10000 bootstrap subsamples (size = 80 % of neurons) and evaluated *H* for the two feature spaces independently.

### Cluster-similarity analysis across brain regions

We compared the cluster-average activity profiles across brain regions with a pair-wise Pearson correlation analysis. For every cluster 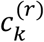 obtained in region *r* we first computed an average dF/F trajectory, 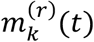 by (1) averaging each neuron’s activity across trials and (2) averaging across all neurons within the cluster. Given two regions, *r*_1_ and *r*_2_, we calculated an 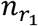 and 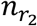 similarity matrix whose (*k, l*) entry was the Pearson correlation between 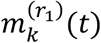 and 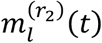, where *n* _*r*_ is the number of clusters in region *r*.

### Forelimb and neuronal rhythmicity analysis

To determine whether MSNs and their cortical and thalamic inputs exhibit rhythmic activity that match the cadence of the contralateral (left) forelimb during locomotion, we followed a three-step procedure encompassing limb tracking, spectral decomposition, and peak-based classification.

#### Video pre-processing and limb trajectory extraction

Video frames were acquired at 30 Hz and analyzed offline with DeepLabCut (DLC) (*52*) to extract the *x* and *y* coordinates of the four limbs. Several DLC models were trained on hundreds of video frames from different animals and recording sessions, to account for variabilities due to animal posture and locomotion idiosyncrasies, light conditions, and limb visibility on the mirror. The four limbs reflected on the mirror were labeled. After automatic labeling, in videos with missing frames, limb coordinates were linearly interpolated based on nearby frames. Labelled values that fell more than four standard deviations outside of the X and Y distributions were likewise replaced with linearly interpolated values.

For the rhythmicity analysis we focused on the left forelimb. Because the forelimb mainly oscillates along a single linear axis related to the direction of movement during locomotion, the two-dimensional trajectory was reduced to a one-dimensional displacement by principal component analysis (PCA): the first principal component (PC_1_) was taken as the scalar limb-position time-series for all subsequent analyses, where the sign of the PC was set so that positive values corresponded to forward motion.

#### Power-spectral density (PSD) estimation

Spectral content of both forelimb trajectory and neural activity was quantified with the Welch’s method (SciPy v1.12.0), using a Hann window of 1024 samples with 50 % segment overlap, which yields a frequency resolution of 0.03 Hz. The limb-position trace and neuronal activity were z-scored at each trial (starting at ladder onset and ending 1 sec after the ladder offset). Trial-wise PSDs were averaged to obtain one spectrum per neuron or per forelimb trajectory.

#### Identification of forelimb-related rhythmic activity

For each session, the averaged limb-position PSD was examined to identify the most prominent spectral peak above 0.03 Hz, yielding the limb’s cadence frequency *f*_*beh*_. The locomotion-related frequency band, defined as the peak’s full width measured at 10 % of its maximum amplitude, served as the reference band for determining forelimb-related neuronal rhythmicity. Within this band, each neuron’s PSD was examined to identify local maxima whose prominence (the height of a peak measured relative to the higher of its two neighboring troughs) exceeded a region-specific threshold that we determined by visual inspection of representative spectra: DLS ≥ 0.80, PF ≥ 1.00, M1 ≥ 0.65 and M2 ≥ 0.65 (PSD units). A neuron was labelled *rhythmic* if its most prominent peak lay inside the band and satisfied the corresponding prominence criterion. Proportions of rhythmic cells were compared between dMSNs and iMSNs, and between projection-defined cortical/thalamic populations, using two-sided Fisher’s exact tests. P-values were Bonferroni-corrected to account for multiple comparisons.

### Phase-tuning analysis

Phase tuning was determined only for neurons deemed to have forelimb-related rhythmic activity identified as above.

#### Stride detection and phase definition

The one-dimensional paw-position trace (PC_1_; see “limb trajectory extraction”) was scanned for successive local minima and maxima. A stride was defined as the interval from one backmost excursion (minimum) to the next, with the intervening frontmost excursion (maximum) marking mid-stride. This interval was mapped onto a phase domain such that the first minimum corresponded to 0° (stride onset), the maximum to 180°, and the subsequent minimum to 360° (stride completion). Phases 0–180° therefore constituted the swing period, when the paw travels forward through the air, whereas phases 180–360° constituted the stance period, when the paw contacts the ladder and moves posteriorly as the body advances relative to the ladder.

#### Stride and neural activity warping

Because individual strides vary in duration, each detected stride was resampled to 360 equally spaced phase points by linear interpolation. The simultaneously recorded neural activity was interpolated onto the same phase grid, yielding one phase-locked activity vector per stride.

### Phase tuning curve

Within each trial, all stride-aligned activity vectors were averaged point-wise to obtain a trial-specific phase-tuning curve, ***T***_**trial**_(***θ***), with ***θ*** ranging from **0**° to 360°. The phase of maximal activity, ***θ***_**peak**,**trial**_ = **argmax**_***θ***_ ***T***_**trial**_(***θ***), was taken as the neuron’s preferred phase on that trial. Preferred phases from all trials of a neuron were combined with a circular mean to give the neuron’s overall preferred phase, ***θ***_**pref**_ = **circmean**{***θ***_**peak**,**trial**_}. We compared preferred-phase distributions between neuron groups using the Watson two-sample ***U***^2^ test, evaluating significance (α = 0.05) with an exact p-value obtained from 10000 random permutations of the group labels for every comparison (*74, 75*).

## Statistics

Statistical tests were selected based on data distributions and significance was set at p < 0.05. Non-parametric tests were used throughout the manuscript where appropriate. No statistical methods were used to pre-determine sample sizes.

## Acknowledgments

We thank David Arakelyan, Ashley Medina, Bobbie Morales, An Wu, and Bin Yu for technical assistance; Lucia Hall and Elanore Hall for assistance with mouse husbandry; Varoth Lilascharoen, Eric Wang, and Jun Hyeok Choi from the Lim laboratory for help with producing AAV helper and rabies viruses; An Wu, Bin Yu, Mikio Aoi and all the Komiyama laboratory members for discussions on this project through the years.

## Author contributions

Conceptualization: EG, HL, TK, RDS, GM

Formal analysis: RDS, SD, JL, GM, HL, EG, TK

Funding acquisition: TK, GM

Investigation: EG, HL, XL, KO, OMA, SWH, HK, SB

Methodology: EG, HL, RDS, GM, BL, TK

Resources: BL, TK, GM

Supervision: TK, GM

Writing – original draft: EG, RDS, GM

Writing – review & editing: EG, RDS, GM, TK and all authors

