## Supplemental Information for "Complementary cortical and thalamic contributions to cell-type-specific striatal activity dynamics during movement"

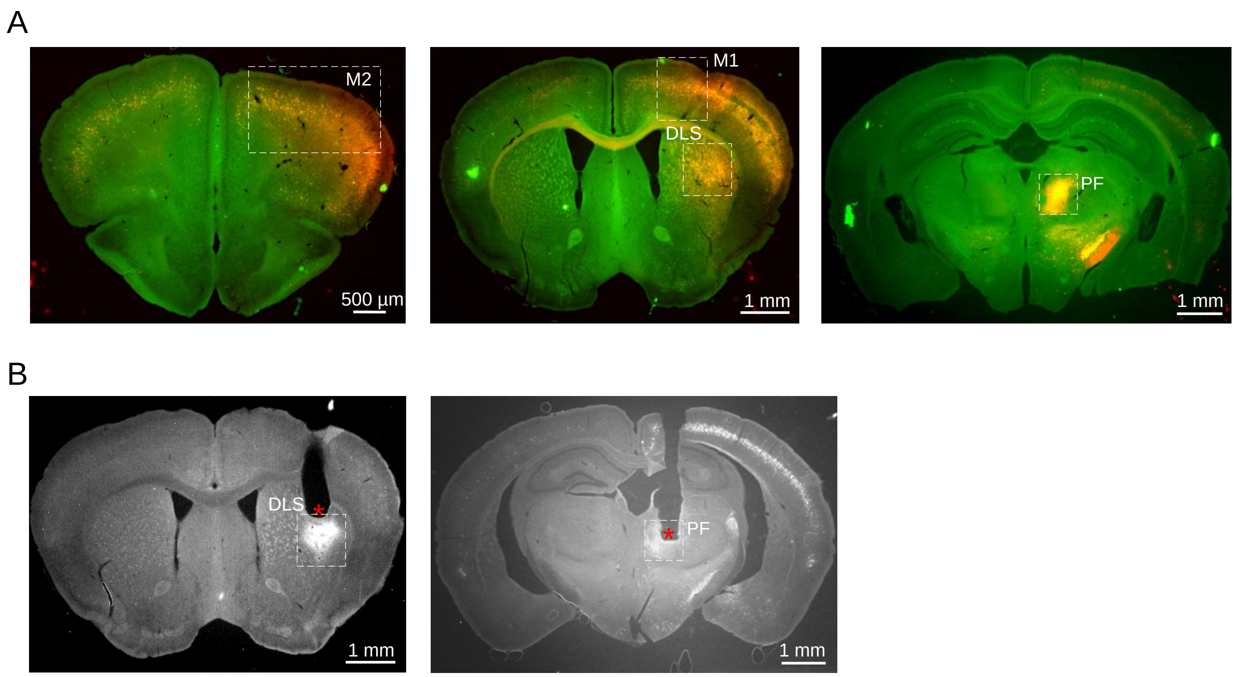


**Fig. S1**

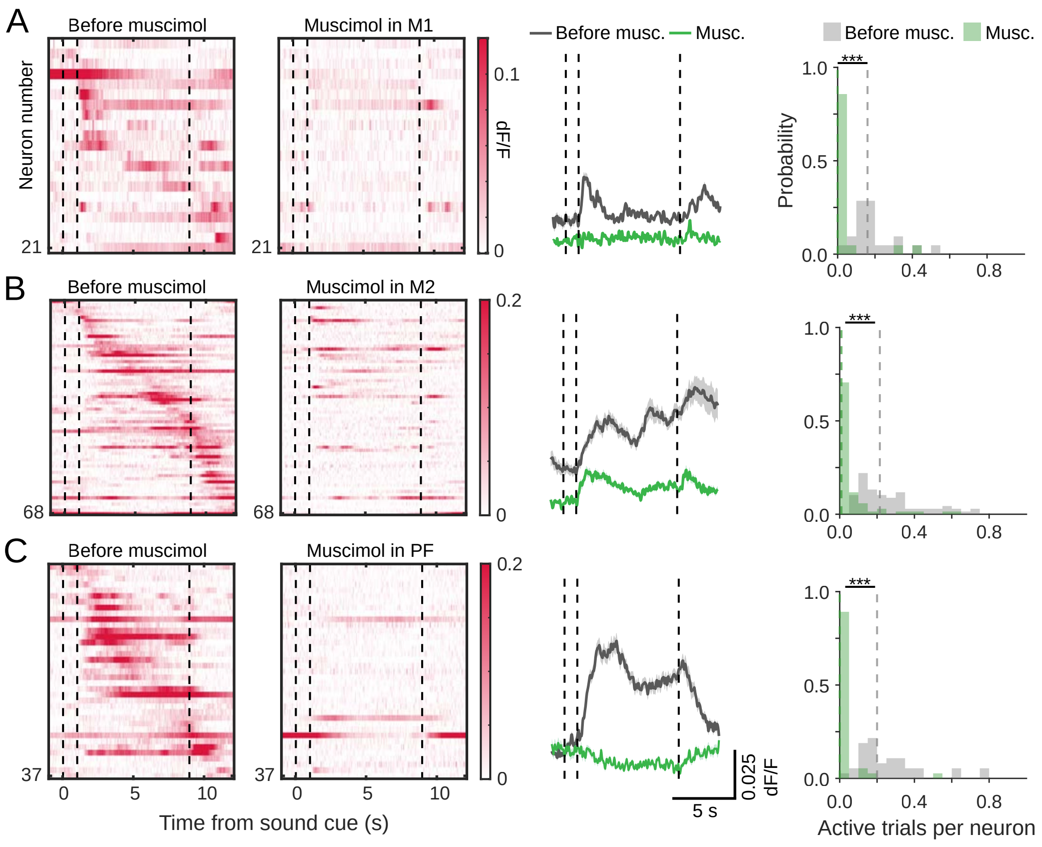
A
B
C
Before muscimol
Muscimol in M1
Muscimol in M2
Muscimol in PF
Neuron number
21
68
37
dF/F
0.1
0.2
0
Before musc.
Musc.
Probability
1.0
0.5
0.0
0.4
0.8
***
0
5
10
Time from sound cue (s)
5 s
0.025 dF/F
Active trials per neuron

**Fig. S2**

| **Model input** | **Classifier model** | **Classification accuracy ± SD (%)** |
|---|---|---|
| Trial-averaged neural activity | Linear SVM | 53.87 ± 5.16 |
| Ensembles of trials | Linear SVM | 52.31 ± 6.53 |
| Ensembles of trials | FCNN with attention | 57.33 ± 3.86 |
| Ensembles of trials | RNN with attention (TEA-net) | 63.21 ± 3.98 |

### Table S1

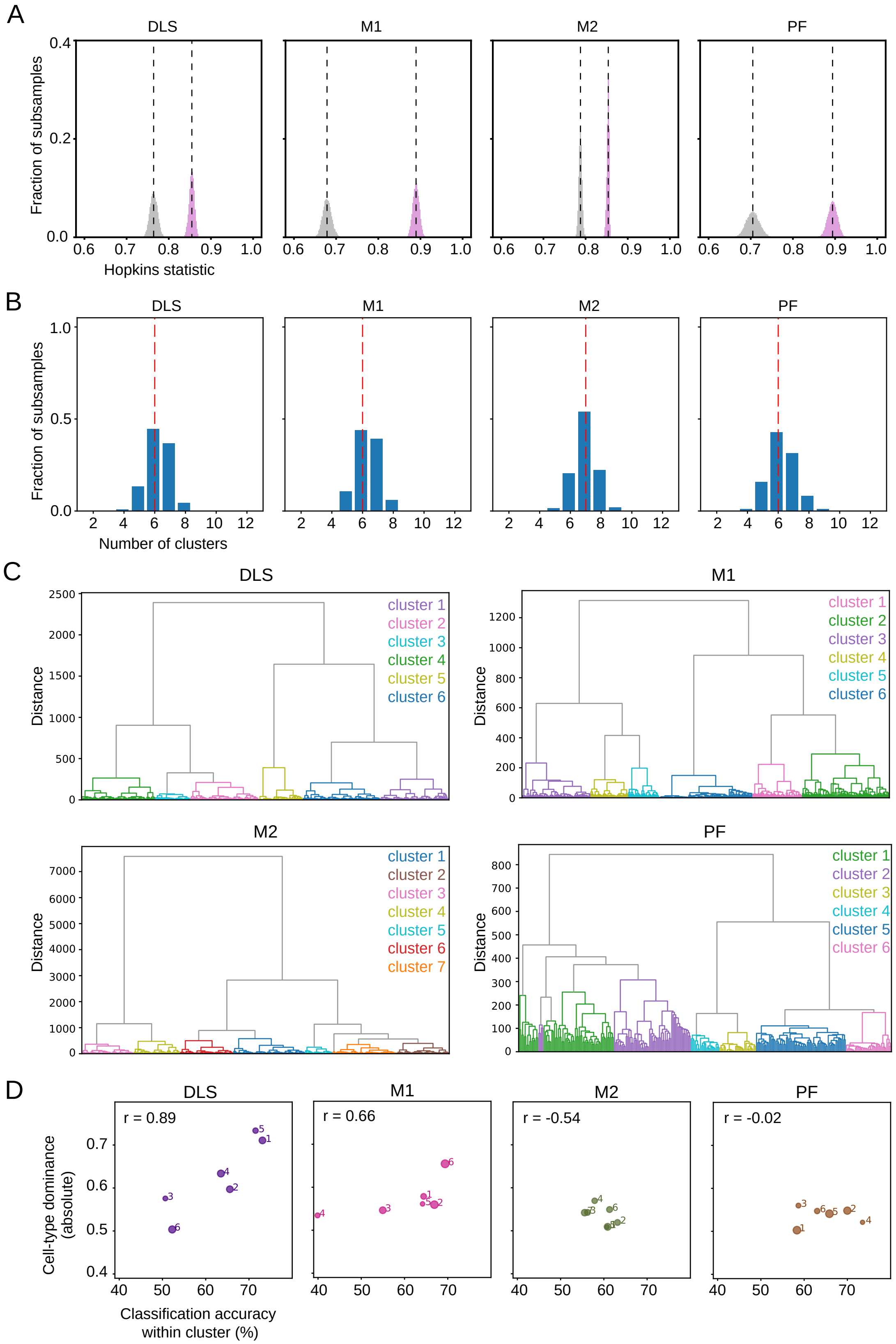

A
DLS
M1
M2
PF
Fraction of subsamples
Hopkins statistic
B
Number of clusters
C
Distance
cluster 1
cluster 2
cluster 3
cluster 4
cluster 5
cluster 6
cluster 7
D
r = 0.89
r = 0.66
r = -0.54
r = -0.02
Cell-type dominance (absolute)
Classification accuracy within cluster (%)


**Fig. S3**

**Fig. S1. Labelling of input neurons in M1, M2 and PF via trans-synaptic spread of pseudo-typed rabies virus and GRIN lens targeting**
**A)** Example coronal brain sections from a Drd1-Cre mouse that show the expression of mRuby-GCaMP6f (mRuby signal in red, GCaMP6f signal in green) via the EnvA-pseudotyped rabies in DLS (injection site) and in the three input regions imaged in this study: M1, M2 and PF. **B)** Example coronal brain sections that show the positioning of the GRIN lens in DLS (left) or in PF (right). Red asterisk indicates the GRIN lens bottom.

**Fig. S2. Muscimol inactivation of cortical or thalamic regions substantially reduces MSN activity**
**A)** From left, trial-averaged activity of MSNs during the ladder task. Dashed lines from left to right in each plot indicate: sound cue onset, ladder onset, and ladder offset; trial-averaged activity of MSNs from the same fields of view, imaged 30 minutes after muscimol injection into M1; average activity of all DLS neurons before and after muscimol injection (mean ± SEM); fraction of active trials in the imaged neurons before and after muscimol injection (n = 21 neurons, of which 14 dMNSs from n = 1 mouse and 7 iMSNs from n = 1 mouse). **B)** Same as A) but for muscimol injections in M2 (n = 68 neurons, of which 24 dMSNs from 3 mice and 44 iMSNs from 4 mice). **C)** Same as A-B) but for muscimol injections in PF (n = 37 neurons, of which 12 dMSNs from 2 mice and 25 iMSNs from 3 mice). Dotted lines indicate medians. Paired Wilcoxon rank-sum test, $p < 0.001$ for all comparisons after vs. before manipulation.

**Table S1. Comparison of cell-type classification performance of TEA-net vs. alternative models**
Classification accuracy of dMSNs and iMSNs obtained by training different classifier models with trial-averaged neural activity or random ensembles of single trials. Classification accuracy was highest for TEA-net. Wilcoxon rank-sum test, TEA-net vs. SVM (trial-averaged data) $p = 0.004$; TEA-net vs. SVM (ensemble data) $p < 0.001$; TEA-net vs. FCNN $p = 0.011$. SVM: Support Vector Machine, FCNN: fully connected neural network, RNN: recurrent neural network, TEA-net: Trial Ensemble Attention Network.

**Fig. S3. Clusterability, optimal number of clusters and hierarchical-clustering dendrograms based on the latent representations for each region.**
**A)** Hopkins statistic distributions of trial-averaged neural activity (in gray) and of the latent space representations (in violet) obtained from 10000 random subsamplings of 80% of the neurons in DLS, M1, M2 and PF. The latent representations yield consistently higher Hopkins values than the corresponding neural activity traces across all four regions, indicating that the latent space possesses a greater likelihood to form clusters. Hopkins statistic for latent representation and neural activity in each region, respectively (mean ± SD): DLS, 0.85 ± 0.01, 0.76 ± 0.01; M1: 0.89 ± 0.01, 0.68 ± 0.01; M2: 0.85 ± 0.002, 0.79 ± 0.003; PF: 0.89 ± 0.01, 0.71 ± 0.02. Dotted lines indicate the median of the distributions. **B)** Distributions of the number of clusters for the latent space representations, obtained from 10000 random subsamplings of 80% of the neurons. The mode of the distributions coincides with the number of clusters obtained when applying the Dynamic Tree cut approach to the complete datasets in DLS, M1, M2 and PF (red dotted line). **C)**

Dendrograms obtained from the hierarchical clustering of the latent space representations of DLS, M1, M2 and PF neurons. Colors indicate the different clusters of neurons obtained in each region (see Fig. 3). **D)** Correlation coefficients between cell-type dominance and classification accuracy in each cluster, for each region. Circle size in each plot corresponds to the cluster size.
